# Synthetic peptide-induced internalization of biomolecules into various plant and algal cells via micropinocytosis

**DOI:** 10.1101/630301

**Authors:** Jo-Ann Chuah, Masaki Odahara, Yutaka Kodama, Takaaki Miyamoto, Kousuke Tsuchiya, Yoko Motoda, Takanori Kigawa, Keiji Numata

**Affiliations:** Biomacromolecules Research Team, RIKEN Center for Sustainable Resource Science, Saitama 351-0198, Japan.; Center for Bioscience Research and Education, Utsunomiya University, Tochigi 321-8505, Japan.; Laboratory for Cellular Structural Biology, RIKEN Center for Biosystems Dynamics Research, Yokohama 230-0045, Japan.

## Abstract

Efficient intracellular delivery of biomolecules is important for many different biological and biotechnological applications in living organisms, and is a prerequisite for certain types of fundamental and applied research. One major challenge is the delivery of unmodified, functional cargoes in a simple, time-efficient, and high-throughput manner. Herein, we present an efficient strategy that uses fusion peptides containing cell penetrating peptide, endosomal escape domain, and a sarcosine linker to introduce biomolecules, namely fluorescent protein and dextran, via macropinocytosis into the cells of various land plants and microalgae. Our peptide-mediated delivery system allows for high-throughput delivery of functional biomolecules within a few minutes to a few hours as well as open new possibilities for biology and biotechnology using difficult-to-transfect cell types.

## Introduction

The ability to introduce biomolecules, such as nucleic acids, proteins, sugars, and small molecules, into living cells is a prerequisite for certain types of fundamental and applied research (Kwak et al., 2019; Torney et al., 2007; Whitehead et al., 2014). Many plants and algae are used as important model systems to elucidate key biological processes; however, effective intracellular delivery of biomolecules, especially proteins, has been a road block that prevents numerous cell-based experiments and high-throughput screens. This is because ribonucleoproteins for genome editing are used for various experiments in biology and biotechnology fields (Woo et al., 2015; Yin et al., 2017). To introduce biomolecules into cells, transfection methods using electroporation or polyethylene glycol with protoplast preparation are generally used; however, these methods are cumbersome, time-consuming, and often yield inconsistent results (Birch, 1997). Microinjection and particle-based methods require expensive instruments and are inherently low-throughput (Newell, 2000). In contrast to these methods, cell-penetrating peptides (CPPs) are a molecule delivery tool that has the intrinsic ability to translocate a diverse range of macromolecules across the plasma membrane in their biologically active form, albeit more widely used for mammalian than plant cells thus far (Chugh et al., 2010; Numata et al., 2018; Numata and Kaplan, 2010). The first CPP, Tat (Frankel and Pabo, 1988; Green and Loewenstein, 1988), was identified in 1988 and delivers macromolecules into cells by first stimulating its own uptake through the induction of macropinocytosis, a specialized form of endocytosis, followed by endosomal escape (Wadia et al., 2004). A significant enhancement in cytoplasmic delivery using Tat was recently reported by Lönn and co-workers through the incorporation of a polyethylene glycol linker and multiple types of synthetic endosomal escape domain (EED) (Lonn et al., 2016).

To develop an efficient strategy for biomolecules delivery into plant cells, we examined Tat-derived peptides comprising the functional region (amino acid residues 49‒57, RRRQRRKKR) (Vives et al., 1997), and employed the retro-inverso (D-) enantiomer of Tat (dTat), which has been shown to provide the peptide with longer-lasting biological activities, presumably because of its resistance against proteases (Schorderet et al., 2005), and a marked enhancement (>100-fold) in the cell internalization rate (Wender et al., 2000). In addition, we selected EEDs optimized for several human cell lines (Lonn et al., 2016), namely, two types of EEDs with different hydrophobic motifs: the first (EED4) containing two aromatic indole rings (-GWWG), and the second (EED5) containing one indole ring and two aromatic phenyl groups (-GFWFG) (Fig. 1A). To avoid cytotoxicity of dTat and EEDs, which was mentioned in a previous study (Lonn et al., 2016) as well as to maintain their original secondary structures, we included a sarcosine-based molecular spacer between the dTat domain and the EED. Six residues of sarcosine has high molecular flexibility, water solubility, and long-circulating properties with negligible nonspecific adsorption *in vitro* and *in vivo* (Weber et al., 2016). In this study, the designed peptides, dTat-Sar-EED4 and dTat-Sar-EED5, were evaluated as synthetic peptides to internalize proteins and dextran into plant suspension cells, intact land plant cells, and algae, which are well known as difficult-to-transfect cell types.

**Figure 1.**
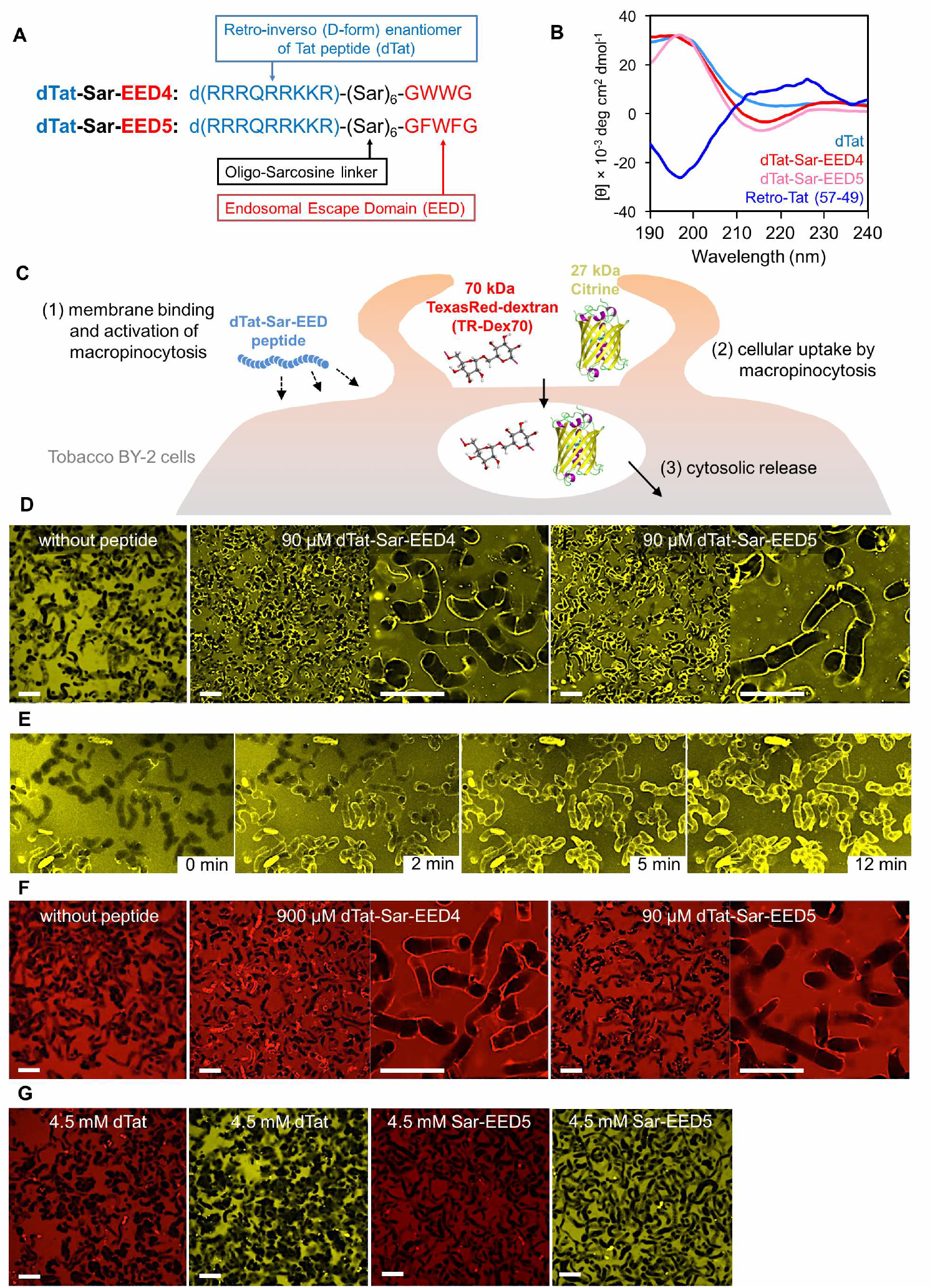
Cargo internalization into tobacco BY-2 cells mediated by the dTat-Sar-EED peptides. (**A**) Amino acid sequences and functions of the individual domains constituting the dTat-Sar-EED peptides. (**B**) Circular dichroism spectra of dTat-Sar-EED peptides as well as dTat and Retro-Tat (57-49) peptides as controls. (**C**) Schematic representation of dextran and Citrine delivery into tobacco BY-2 cells through the induction of macropinocytosis by dTat-Sar-EED peptides. (**D**) Confocal microscopic images showing cells treated for 1 h with Citrine alone, and in combination with each dTat-Sar-EED peptide at the minimal effective concentrations. Scale bars indicate 50 µm. (**E**) Time-lapse images of Citrine internalization by 90 µM of dTat-Sar-EED4. (**F**) Confocal microscopic images showing cells treated for 1 h with dextran (TR-Dex70) alone, and in combination with each dTat-Sar-EED peptide at the minimal effective concentrations. Scale bars indicate 50 µm. (**G**) Confocal microscopic images showing TR-Dex70 (red) and Citrine (yellow) internalization into tobacco BY-2 cells mediated by dTat (left) or Sar-EED5 (right) as controls. The cells were treated for 1 h with TR-Dex70 or Citrine in combination with dTat or Sar-EED5 at 4.5 mM. Scale bars indicate 50 µm.

## Results and Discussion

### Secondary structures of the peptides

The designed peptides, dTat-Sar-EED4 and dTat-Sar-EED5, were easily synthesized by the traditional solid-phase synthesis method without the need for special chemical reactions (Fields and Noble, 1990). Circular dichroism analysis was carried out to analyze and compare the secondary structures of the dTat-Sar-EED peptides as well as the original dTat and Retro-Tat (57-49) as controls (Figure 1B). Both dTat-Sar-EED4 and dTat-Sar-EED5 lack a defined secondary structure and their spectra are mirror images of Retro-Tat’s spectrum, which is typical of that for unstructured peptides with an absorbance minimum at 195 nm (Yang et al., 1986). dTat showed a similar structure to the dTat-Sar-EED peptides, indicating that the sarcosine linker prevents the hydrophobicity of the EED from affecting the secondary structure of the dTat domain.

### Protein internalization into plant suspension cells

The dTat-Sar-EED peptides were evaluated for their ability to promote the intracellular uptake of the 27 kDa fluorescent protein Citrine into tobacco (*Nicotiana tabacum*) Bright Yellow-2 (BY-2) suspension cells (Figure 1C). In the presence of Citrine, dTat-Sar-EED4 was administered to the cells at a wide concentration range of 9 nM to 9 mM (**Figure S1**). Confocal microscopic analysis revealed that the cells could internalize Citrine within a short incubation duration of 1 h with 90 µM and higher concentrations of both dTat-Sar-EED peptides (Figure 1D). Citrine fluorescence was not seen in cells in the absence of the peptide, whereas when the peptide was included, distinct fluorescent signals of internalized Citrine were visible in the periphery of most cells and were detected in the nucleus (**Figure S1**). We carried out a time-lapse experiment to observe cellular internalization of Citrine (Figure 1E). Remarkably, internalization of Citrine occurred rapidly using either dTat-Sar-EED4 or dTat-Sar-EED5 within approximately 2 to 12 minutes following peptide addition, making this a high-throughput as well as time-efficient process (**movie S1 and S2**).

### Cytotoxicity of dTat-Sar-EED peptides

Exposure of BY-2 cells to higher peptide concentrations (900 µM to 9 mM) for longer durations (24‒48 h) resulted in cytotoxicity, as evident from the cytoplasmic shrinkage and detachment from the cell wall observed in the confocal microscopic analyses (**Figure S2A**). Thus, we evaluated the viability of BY-2 cells at effective peptide concentrations and incubation durations (**Figure S2B**). Cells that were untreated (1‒48 h) were included in the analysis for comparison. Notably, dTat-Sar-EED5 affected cell viability (50‒70% cell death) to a greater extent than dTat-Sar-EED4 (20‒40% cell death) at all effective concentrations, possibly owing to the two constituent phenylalanine residues. dTat-Sar-EED5 was far more toxic against BY-2 cells than dTat-Sar-EED4 at an equal peptide concentration needed for Citrine and dextran delivery, making dTat-Sar-EED4 more feasible for use with tobacco BY-2 cells. At higher peptide concentrations (900 µM to 9 mM) for longer durations (24‒48 h), both dTat-Sar-EED peptides demonstrated cytotoxicity to BY-2 cells. To reduce such cytotoxic effects, we screened peptide concentrations between the ineffective dose (9 µM) and the minimal effective dose (90 µM) for Citrine internalization; however, decreased peptide amounts were accompanied by a decline in Citrine uptake into cells. The ability of Citrine-internalized cells to proliferate and cell viability was monitored over a 10-day duration (**Figure S2C**). Two representative dTat-Sar-EED4 concentrations were selected (both effective, 90 and 900 µM) and untreated cells as well as cells treated with Citrine alone were included for comparison. While the higher peptide concentration (900 µM) inhibited cell growth and resulted in a higher percentage of dead cells (stained by Evans blue), it is noteworthy that the growth rate and viability of cells treated with the optimal peptide concentration (90 µM) in the presence of Citrine were similar with that of untreated and Citrine-only controls.

### Dextran internalization into BY-2 cells

The delivery of another biomolecule, Texas Red-labeled 70 kDa dextran (TR-Dex70), was subsequently examined under the same conditions (Figure 1F and **S3**). High levels of fluorescence of internalized TR-Dex70 could be seen surrounding the peptide-treated cells, but fluorescence was completely absent in cells exposed to TR-Dex70 without peptide, regardless of the duration of incubation. Lower concentrations of dTat-Sar-EED4 (90–900 µM) enabled the intracellular uptake of dextran with prolonged incubation of up to 24 h (**Figure S3B**), while extending the incubation duration up to 48 h did not improve TR-Dex70 internalization (**Figure S3C**). For comparison, dTat-Sar-EED5 was administered to the cells at the same concentration range (9 nM to 9 mM) in the presence of TR-Dex70 (**Figure S3D**) and the results showed that 90 µM of dTat-Sar-EED5 and a 1-h duration were sufficient for dextran delivery into BY-2 cells.

### Importance of fusion design

We considered the possibility of the individual domains dTat, Sar-EED4, or Sar-EED5 having similar capacities for dextran and Citrine delivery into cells as that of dTat-Sar-EED4 and dTat-Sar-EED5. Based on experiments using the individual domains, no distinguishable fluorescent signals could be observed in the cells at relatively higher peptide concentration, 4.5 mM (Figure 1G). Via a time-lapse observation of cellular internalization, we found that Citrine was not internalized at all using 90 µM dTat under identical experimental conditions (**movie S3**). This implies that exogenously added biomolecules need to enter the cells, which is facilitated by the dTat domain, and be adequately released into the cytosol, which is assisted by the EED. While it is expected that neither of the EEDs possess cell-penetrating functions, the lack of fluorescence surrounding cells treated with dTat suggests that although dTat is an integral competent in intracellular translocation, its efficiency is not sufficient to transport the cargo molecules across the cell wall, plasma membrane, and endosomes into the cytosol.

### Internalization pathway

To confirm the dTat-Sar-EED peptide-induced pathways such as macropinocytosis for cellular uptake of Citrine and TR-Dex70, we analyzed the internalization pathways via inhibitor assays (Figure 2A). We incubated cells with a fluorescent fluid-phase macropinocytosis marker, neutral TR-Dex70 (Araki et al., 1996; Oliver et al., 1984), in combination with dTat-Sar-EED4 or dTat-Sar-EED5. Cells were also treated with various types of endocytosis inhibitors or incubated at 4°C to investigate the involvement of other internalization pathways (Figure 2B). The neutral TR-Dex70 marker was primarily taken up by macropinocytosis, based on the almost complete suppression of fluorescence intensity in the presence of amiloride (EIPA) (Commisso et al., 2013) and cytochalasin D (CytD) (Nakase et al., 2004). Very slight inhibition was observed with chlorpromazine (CPZ) (Wang et al., 1993) and filipin (Schnitzer et al., 1994). Based on the fluorescence intensity, we quantify the inhibition behaviors (Figure 2C). The findings were consistent for both dTat-Sar-EED peptides; signifying negligible involvement of clathrin-or caveolae-mediated endocytosis. Energy-independent pathways do not contribute to cargo internalization as cells were unable to internalize dextran at 4°C.

**Figure 2.**
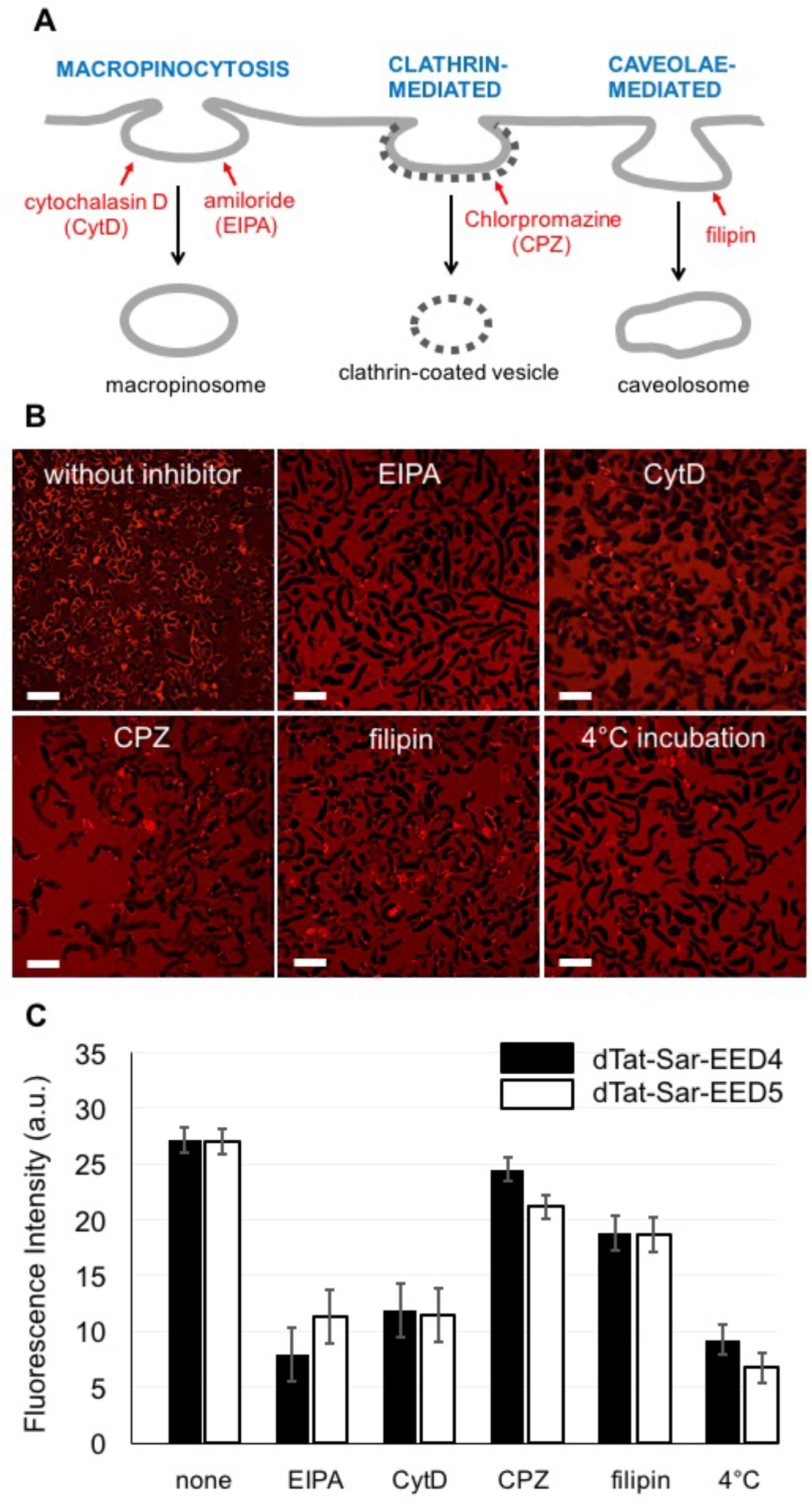
Effect of various inhibitors on the cellular uptake of dextran (TR-Dex70). (**A**) Schematic representation of the different endocytic internalization pathways and corresponding inhibitors. (**B**) Confocal microscopic images showing TR-Dex70 internalization into BY-2 cells using dTat-Sar-EED5 without and with the presence of the inhibitors amiloride (EIPA), cytochalasin D (CytD), chlorpromazine (CPZ), and filipin. Scale bars indicate 50 µm. (C) The bar graph denotes the quantification of fluorescence intensity corresponding to the images in dextran-internalized cells. Data represent the mean values ± s.d. (*n* = 6).

### Internalization into intact land plants

To further explore the applicability of dTat-Sar-EED peptides, we used similar conditions to deliver the fluorescent molecules into intact leaves of *Arabidopsis thaliana*, a model system for land plants. Distinct localization of Citrine in the cytosol was seen when dTat-Sar-EED4 or dTat-Sar-EED5 was present, whereas Citrine could not be internalized without a peptide (Figure 3). In this case, 90 µM of either peptide was inefficient in translocating Citrine into the cytosol, and hence a higher concentration of peptide (900 µM) was needed for Citrine localization in the cytosolic compartment of epidermal and mesophyll cells. Likewise, TR-Dex70 was observed only in the intracellular space of cells without inclusion of dTat-Sar-EED peptides (**Figure S4**). In contrast, dTat-Sar-EED4 and dTat-Sar-EED5 (90 µM for both peptides) enabled cellular internalization of TR-Dex70 that uniformly filled the cytosolic and vacuolar compartments, evident in both epidermal as well as mesophyll layers of the leaf. Neither dTat nor Sar-EEDs alone could transport TR-Dex70 or Citrine across the intracellular space of cells into the cytosol (Figure 3 and **S4**). Notably, similar to Arabidopsis, dTat-Sar-EED4 successfully delivered Citrine into another land plant, the model liverwort *Marchantia polymorpha* (Figure 4). *M. polymorpha* belongs to the bryophytes, the basal group of all land plants, and is known hardly internalized with exogenous proteins (Bowman et al., 2016). When we applied dTat-Sar-EED4 with Citrine to the germinated spores of *M. polymorpha*, we observed internalization of Citrine. Given that protein delivery by dTat-Sar-EED4 could be used in both the basal land plant *M. polymorpha* and the angiosperm Arabidopsis, the delivery system may be applicable in a wide variety of land plants.

**Figure 3.**
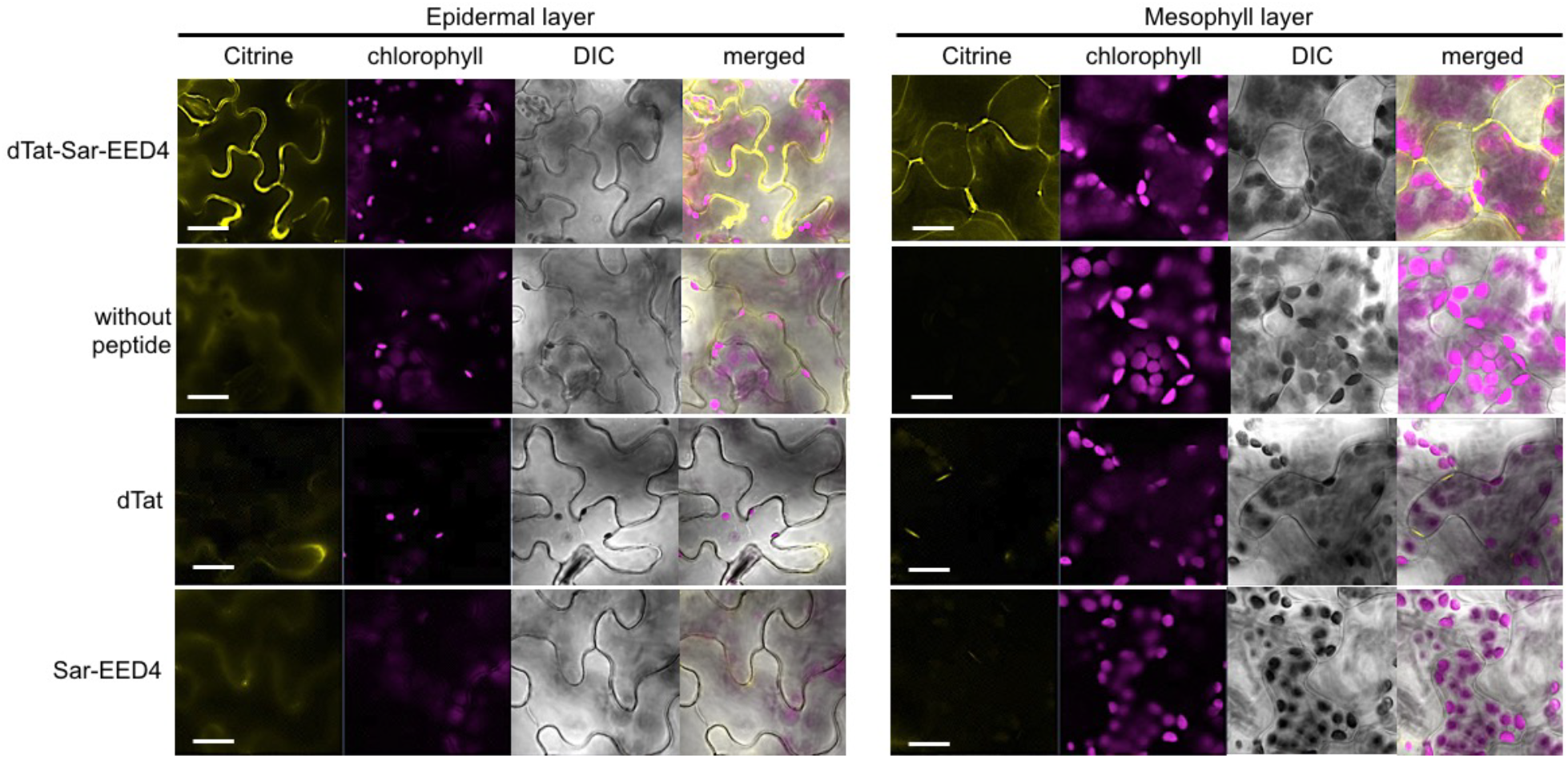
Citrine internalization into *Arabidopsis thaliana* leaf epidermal and mesophyll cells mediated by the dTat-Sar-EED4. Confocal microscopic images showing Arabidopsis leaf cells treated for 1 h with Citrine alone and in combination with each dTat-Sar-EED4, dTat and Sar-EED4 at a concentration of 900 µM. Scale bars indicate 10 µm.

**Figure 4.**
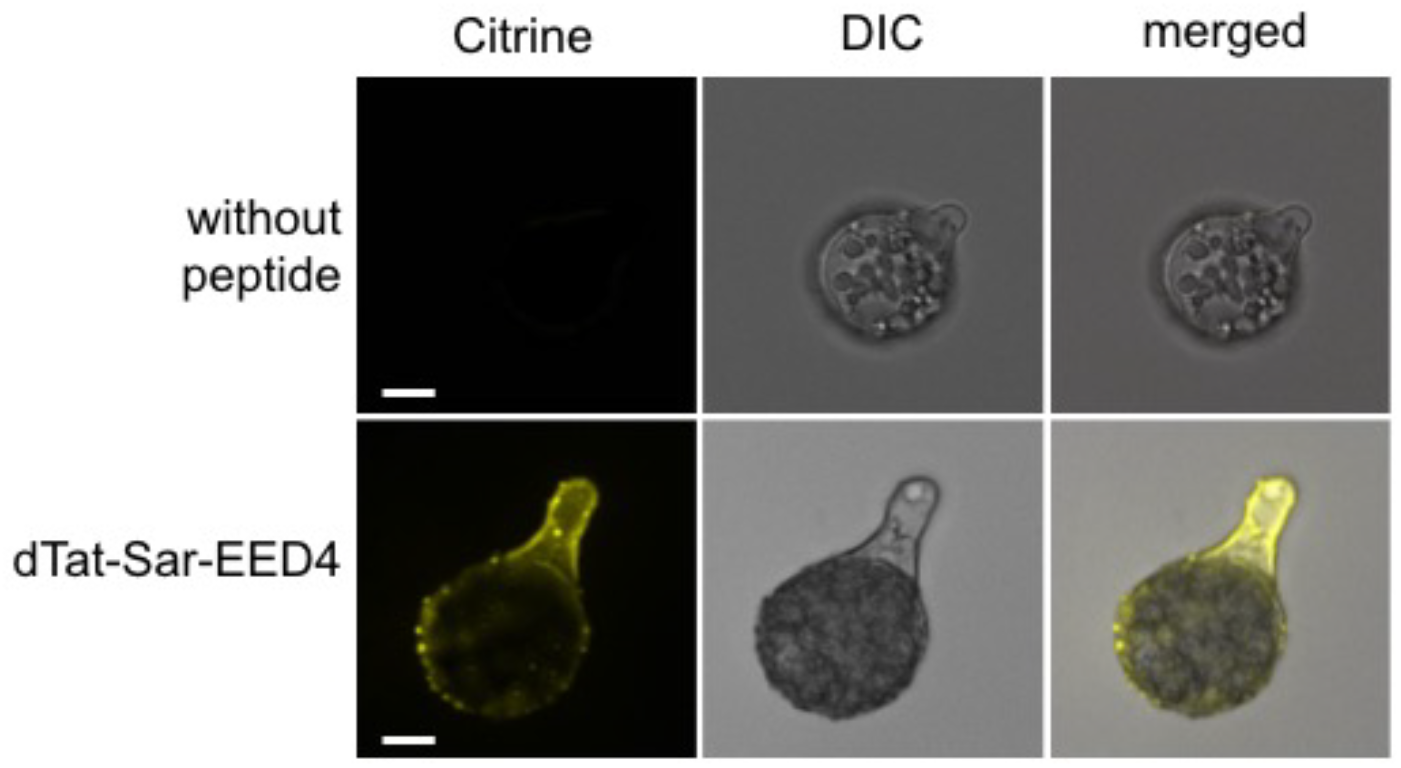
Citrine internalization into cells of *Marchantia polymorpha* mediated by dTat-Sar-EED4. Confocal microscopic images showing the germinated spore from a sporangia of the liverwort *M. polymorpha* after treatment with Citrine and the dTat-Sar-EED4 peptide for 1 h. Scale bars indicate 10 µm.

### Applicability to microalgae

We then tested the functionality of the dTat-Sar-EED peptide-mediated delivery system on unicellular eukaryotes, the microalgae *Chlamydomonas reinhardtii* and *Euglena gracilis*. TR-Dex70 was easily internalized with only 9 µM dTat-Sar-EED4 or dTat-Sar-EED5 by *C. reinhardtii* cells within 3 h, whereas Citrine was only internalized with 900 µM of dTat-Sar-EED5 (Figure 5 and **S5**). According to a previous report on cell-penetrating peptides and Chlamydomonas, *C. reinhardtii* cells seem to need endocytic pathways in addition to macropinocytosis-mediated protein internalization (Kang et al., 2017). Therefore, fewer endocytic pathways induced by dTat-Sar-EED5, which is a strong macropinocytosis inducer for *C. reinhardtii* cells due to its aromatic phenyl groups, would lead to relatively low cellular uptake of Citrine.

**Figure 5.**
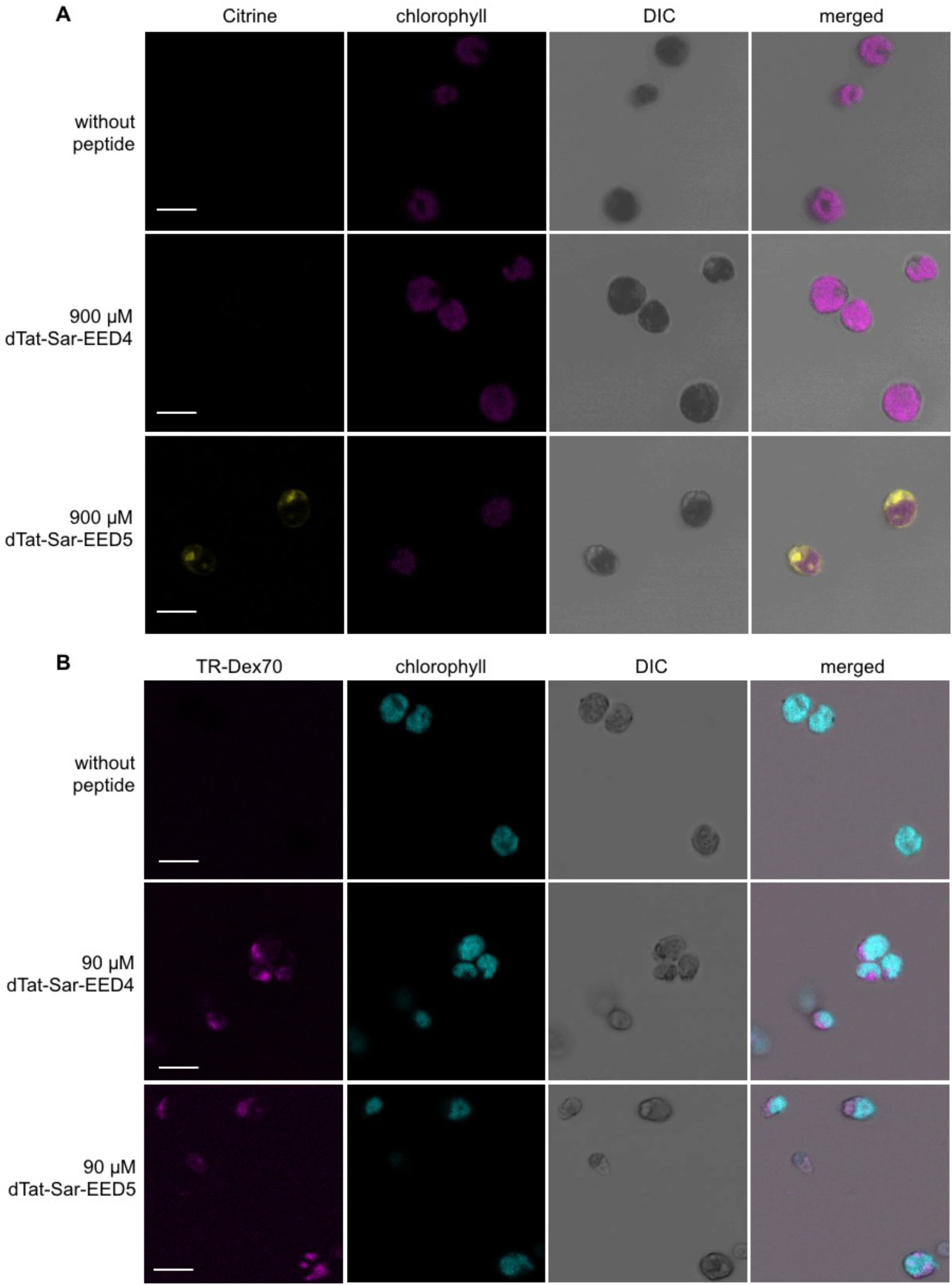
Citrine and dextran (TR-Dex70) internalization into *Chlamydomonas reinhardtii* cells mediated by the dTat-Sar-EED peptides. (**A**) Confocal microscopic images showing *C. reinhardtii* cells treated for 3 h with Citrine alone and in combination with each dTat-Sar-EED peptide at a concentration of 900 µM. Scale bars indicate 10 µm. (**B**) Confocal microscopic images showing *C. reinhardtii* cells treated for 3 h with TR-Dex70 alone and in combination with each dTat-Sar-EED peptide at a concentration of 90 µM. Scale bars indicate 10 µm.

For *E. gracilis*, one of poorly transformable organisms with proteins, Citrine was able to be introduced into cells with 9 nM dTat-Sar-EED4 (Figure 6A). The efficiencies of both dTat-Sar-EEDs for Citrine delivery were equivalent, requiring 9 nM and higher concentrations of peptide and an incubation duration of 3 h or more (**Figure S6A**). In the case of TR-Dex70, no noticeable changes were seen in the cellular uptake efficiency of TR-Dex70 at 6 h post-treatment, while at 24 h post-treatment, cells appeared to have internalized TR-Dex70 with 9 nM of peptide (Figure 6B and **S6B**). Intriguingly, dTat-Sar-EED5 functions more efficiently in *E. gracilis* cells than in tobacco BY-2 cells; a minimum concentration of 9 nM peptide enabled delivery of TR-Dex70 into *E. gracilis* cells within 3‒24 h. We reasoned that the difference in hydrophobic motifs between the two EEDs, i.e., the additional phenylalanine residues in EED5, that was toxic to the BY-2 cells may instead provide a beneficial interaction with the pellicle (Sommer, 1965) (outer layer of protein-rich soft tissue) of *E. gracilis* cells, enabling easier access into the interior.

**Figure 6.**
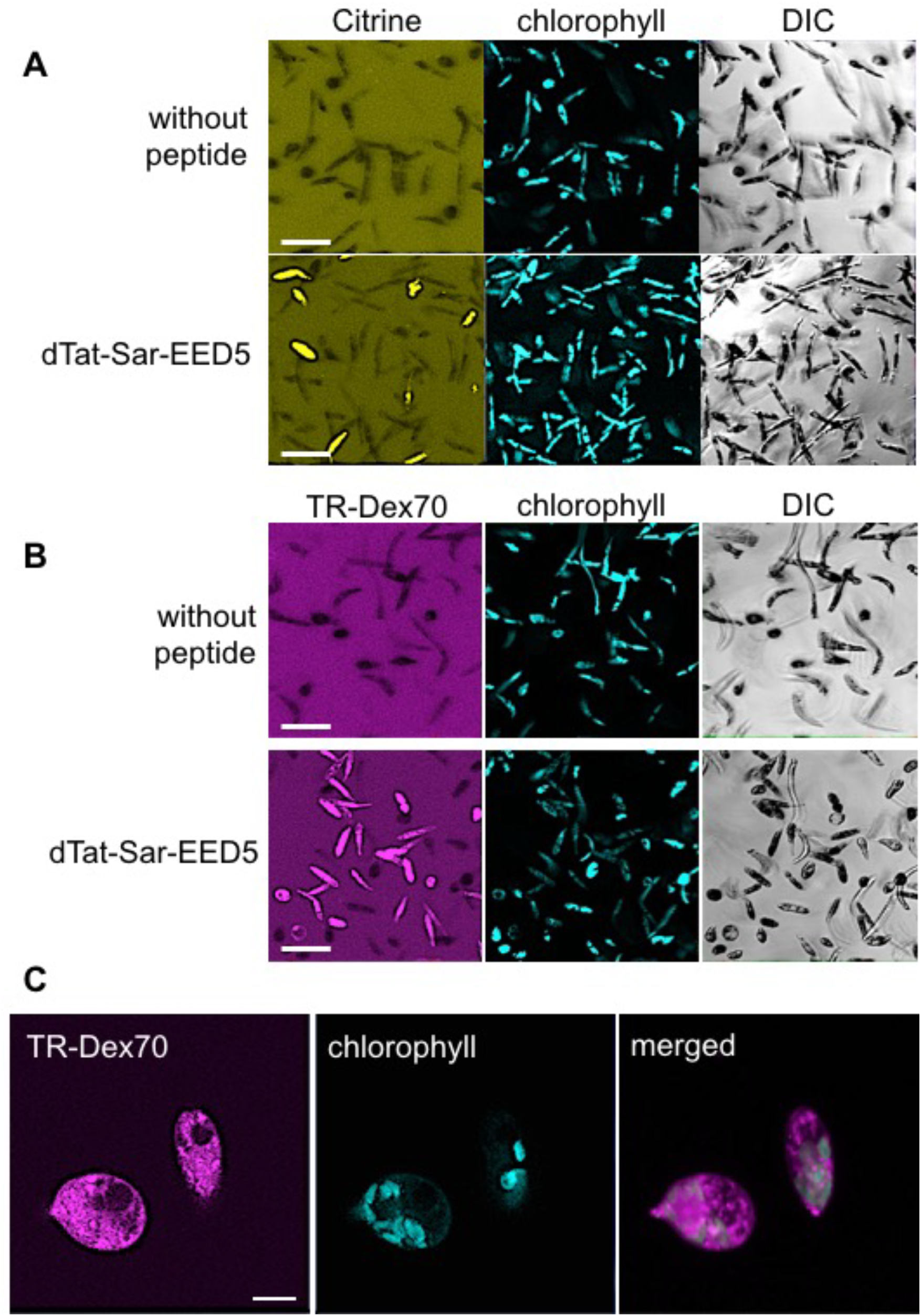
Citrine and dextran (TR-Dex70) internalization into *Euglena gracilis* cells mediated by the dTat-Sar-EED5. Confocal microscopic images showing *E. gracilis* cells treated for 6 h with Citrine (**A**) or TR-Dex70 (**B**) alone and in combination with dTat-Sar-EED5 at a concentration of 90 nM. Scale bars indicate 50 µm. (**C**) Enlarged image of TR-Dex70-internalized *E. gracilis* cells. Scale bars indicate 10 µm.

### Potential applications of dTat-Sar-EED peptides

In this study, we introduced a simple strategy to deliver biomolecules into diverse cell types ranging from cultured tobacco BY-2 cells, to the intact land plants *A*. *thaliana* and *M. polymorpha*, to the microalgae *C. reinhardtii* and *E. gracilis.* The exogenously added macromolecule (70 kDa dextran) and fluorescent protein (27 kDa Citrine) could efficiently cross the cell membrane in an energy-dependent manner, by virtue of the dTat-Sar-EED peptides, and localize to the cytosol as well as the nucleus in some cases. Cargo transduction facilitated by these peptides occurs by macropinocytosis, allowing for high-throughput delivery of target biomolecules, and the process is also rapid, occurring on time-scales ranging from a few minutes to a few hours. Our discovery of this versatile method, which can be applied to deliver a number of biologically-active macromolecular cargos, such as ribonucleoproteins for genome editing, may open new possibilities for research and technological development using difficult-to-transfect cell types.

## Experimental procedures

### Materials

dTat-Sar-EED4 [d(RRRQRRKKR)-(Sar)_6_-GWWG, 2360.69 Da], dTat-Sar-EED5 [d(RRRQRRKKR)-(Sar)_6_-GFWFG, 2468.83 Da], dTat [d(RRRQRRKKR), 1339.62 Da], Sar-EED4 [(Sar)_6_-GWWG, 1039.09 Da], and Sar-EED5 [(Sar)_6_-GFWFG, 1147.23 Da] were synthesized by the Research Resources Center of RIKEN Brain Science Institute. Retro-Tat (57‒49) [RRRQRRKKR, 1339.62 Da] was synthesized by Eurofins Genomics LLC. Citrine (27 kDa) was synthesized and purified as described previously(Ng et al., 2016). Texas Red-labeled dextran (TR-Dex70, 70 kDa) was purchased from Invitrogen (Carlsbad, CA). Evans Blue, amiloride (EIPA), cytochalasin D, chlorpromazine, and filipin were purchased from Sigma-Aldrich (St. Louis, MO).

### Culture and growth conditions of tobacco BY-2 cells

Tobacco (*Nicotiana tabacum*) BY-2 cell suspension cultures were purchased from RIKEN BioResource Center. The cells were maintained in a modified Linsmaier and Skoog medium in the dark at 26°C, 130 rpm, and subcultured at one-week intervals as described previously (Nagata et al., 1992).

### Circular dichroism (CD) spectroscopy

The CD spectra of peptides (10 μM) in water were acquired using a Jasco J-820 CD spectropolarimeter. Background scans were obtained for water. Measurements were made using a quartz cuvette with a 0.1-cm pathlength. Each spectrum represents the average of ten scans from 190 to 240 nm with a 1-nm resolution, obtained at 200 nm/min with a bandwidth of 1 nm.

### Peptide transduction for tobacco BY-2 cells

Cargo internalization into tobacco BY-2 cells was performed in 96-well microplates. Exponentially growing cells (3 days after subculture) were diluted to an OD_600_ of 0.5 with culture medium and 80 µl was added to each well. Cells were then treated with 9 nM to 9 mM of peptide in the presence of 100 µg/ml of TR-Dex70 or Citrine. Culture medium was added to each well to a final volume of 100 µl followed by incubation at 26°C, 130 rpm, for 1‒48 h before analysis. To study the effect of inhibitors on the cellular uptake of dextran, cells were pretreated with various inhibitors (1 mM amiloride, 10 µM cytochalasin D, 10 µg/ml chlorpromazine, and 3 µg/ml filipin) at 26°C, or preincubated at 4°C for 2 h, followed by the addition of dextran and peptide (900 µM of dTat-Sar-EED4 or 90 µM of dTat-Sar-EED5) as described above. For quantification, 100 µl of dextran-internalized cells was washed three times by centrifugation (200 × *g*, 10 min) followed by resuspension in Complete Minimal (CM) medium. Cells were then transferred onto 96-well microplates and fluorescence intensity was determined using excitation and emission wavelengths of 595 nm and 615 nm, respectively.

### Culture conditions and peptide transduction for Arabidopsis, *Marchantia polymorpha*, *Chlamydomonas reinhardtii*, and *Euglena gracilis*

*Arabidopsis thaliana* ecotype Col-0, which serves as a model plant, was grown under the same conditions used previously (Lakshmanan et al., 2013). For experiments using Arabidopsis leaves, a transduction solution consisting of 4.4 mM of peptide, 100 µg/ml of dextran or Citrine, and Milli-Q water to a final volume of 100 µl was prepared. Leaves were then infiltrated with the solution by a syringe as described previously (Lakshmanan et al., 2013) and incubated at 26°C for 1 h before analysis.

Spores of the liverwort *Marchantia polymorpha* were obtained by crossing between the male Tak-1 and female Tak-2 strains (Ogasawara et al., 2013). Spores from sporangia were incubated in 150 µl Milli-Q water for 4 days, and the 4-day-old germinated spores were applied to the transfection assay of Citrine with the dTat-Sar-EED4 peptide. The germinated spores were mixed with dTat-Sar-EED4 (1.7 µM) and Citrine (0.2 mg/ml) and incubated at 22°C for 1 h. The germinated spores were mixed with only Citrine (0.2 mg/ml) as a control.

*Chlamydomonas reinhardtii* wild-type (cc125+) cells were cultivated in Tris Acetate Phosphate (TAP) liquid medium at 23°C under constant light. Cells were stained with TAP medium containing 100 µg/ml of TR-Dex70 or Citrine at variable concentrations of dTat-Sar-EED4 or dTat-Sar-EED5 for 3 and 24 hours.

*Euglena gracilis* cells were maintained in CM medium (pH 3.5) at 26°C, 100 rpm, and subcultured at one-week intervals as described previously (Cramer and Myers, 1952). Cargo internalization into *E. gracilis* cells was performed in 96-well microplates. Cells were cultured to an OD_730_ of 0.23 and 80 µl was added to each well. Cells were then treated with 9 nM to 900 µM of peptide in the presence of 100 µg/ml of TR-Dex70 or Citrine. Culture medium was added to each well to a final volume of 100 µl followed by incubation at 26°C, 100 rpm, for 3‒24 h before analysis.

### Confocal laser scanning microscopy

TR-Dex70 and Citrine internalization into tobacco BY-2, *C. reinhardtii*, and *E. gracilis* cells was directly visualized from the microplate under various magnifications as specified, at excitation wavelengths of 488 nm (for Citrine) and 555 nm (for TR-Dex70) using a confocal microscope (LSM 700/880, Carl Zeiss, Oberkochen, Germany) and Zen 2011 operating software. Time-lapse imaging was performed by capturing 140 frames at 1 msec intervals over a duration of 15 minutes. Arabidopsis leaf samples were prepared and observed as previously described (Lakshmanan et al., 2012). When needed, the cell wall was stained with Calcofluor White solution (0.2 g/l, 10 min) prior to microscopic observation. To observe Citrine fluorescence in *M. polymorpha*, we used a confocal laser scanning microscope SP8X system (Leica Microsystems) with a time-gated method (0.5–12.0 ns) according to a previous report (Kodama, 2016). For excitation, we used a 510-nm laser and collected emissions at 546–566 nm.

### Time-course analysis of cell growth and viability

The growth of untreated, citrine-treated, and citrine-internalized BY-2 cells (with the addition of 90 or 900 µM of peptide) was monitored by performing optical density measurements of the cells at 600 nm using a SpectraMax M2 spectrophotometer (Molecular Devices, Sunnydale, CA) at intermittent time points over a duration of 10 days. Cell viability was determined at corresponding time points by incubating the cells with 0.15 mg/ml of Evans blue in distilled water (1:1), followed by solubilization of bound stain in 50% aqueous methanol containing 1% SDS and spectrophotometric quantification at 600 nm, as detailed in a previous report (Iriti et al., 2006).

### Cell viability assay

BY-2 cells were incubated with various concentrations of the peptides at 26°C for 1‒48 h, and cell viability was evaluated as described above.

### Statistical analysis

SPSS 22.0 for Mac (IBM, Armonk, NY) was employed for statistical analysis. Tukey’s Honest Significant Difference (HSD) test was used for pairwise comparisons among means. Differences between two means were considered statistically significant at *P* < 0.05 and are indicated with asterisks (*). Experimental data are expressed as the means ± standard deviation.

## Acknowledgments

We thank Dr. Takashi Osanai for providing the *Euglena gracilis* strain and Dr. Yoshiki Nishimura for providing the *Chlamydomonas* strain. We also thank Dr. Takayuki Kohchi for providing the *Marchantia polymorpha* strain, and Ms. Mio Hikawa for technical support of the *Marchantia polymorpha* experiment.

## Author contributions

JC and KN conceived the study and designed the experiments. JC performed all the experiments and analyzed the data of BY-2 cells, *A. thaliana*, and *E. gracilis*. MO and YK performed all the experiments and analyzed the data of *C. reinhardtii* and *M. polymorpha*, respectively. TM and KT prepared and chemically characterized the peptides. YM and TK prepared Citrine by cell-free synthesis. JC and KN wrote the manuscript and all authors edited the manuscript. All authors have given approval to the final version of the manuscript.

## Data and materials availability

All data needed to evaluate the conclusions in the paper are present in the paper. Additional data related to this paper may be requested from the authors.

## Supporting Information

Figure S1. Citrine internalization into BY-2 cells mediated by dTat-Sar-EED4 and dTat-Sar-EED5.

Figure S2. Cytotoxicity effect of dTat-Sar-EED peptides on BY-2 cells.

Figure S3. Dextran internalization into tobacco BY-2 cells.

Figure S4. Dextran internalization into *A. thaliana* leaf epidermal and mesophyll cells.

Figure S5. Cargo internalization into *C. reinhardtii* cells.

Figure S6. Cargo internalization into *E. gracilis* cells.

Movie S1. Citrine uptake by tobacco BY-2 cells in the presence of 90 µM dTat-Sar-EED4.

Movie S2. Citrine uptake by tobacco BY-2 cells in the presence of 90 µM dTat-Sar-EED5.

Movie S3. Citrine uptake by tobacco BY-2 cells in the presence of 90 µM dTat as a control.

