## Supporting Information for "Synthetic peptide-induced internalization of biomolecules into various plant and algal cells via micropinocytosis"

Plant Biotechnology Journal
Supporting Information

### Synthetic peptide-induced internalization of biomolecules into various plant and algal cells via micropinocytosis

Jo-Ann Chuah[1], Masaki Odahara[1], Yutaka Kodama[1,2], Takaaki Miyamoto[1], Kousuke Tsuchiya[1], Yoko Motoda[1,3], Takanori Kigawa[3], and Keiji Numata[1]*

[1]Biomacromolecules Research Team, RIKEN Center for Sustainable Resource Science, Saitama 351-0198, Japan.
[2]Center for Bioscience Research and Education, Utsunomiya University, Tochigi 321-8505, Japan.
[3]Laboratory for Cellular Structural Biology, RIKEN Center for Biosystems Dynamics Research, Yokohama 230-0045, Japan.

* Keiji Numata's

**Supplementary Materials**

Figure S1. Citrine internalization into BY-2 cells mediated by dTat-Sar-EED4 and dTat-Sar-EED5.
Figure S2. Cytotoxicity effect of dTat-Sar-EED peptides on BY-2 cells.
Figure S3. Dextran internalization into tobacco BY-2 cells.
Figure S4. Dextran internalization into *A. thaliana* leaf epidermal and mesophyll cells.
Figure S5. Cargo internalization into *C. reinhardtii* cells.
Figure S6. Cargo internalization into *E. gracilis* cells.
Movie S1. Citrine uptake by tobacco BY-2 cells in the presence of 90 μM dTat-Sar-EED4.
Movie S2. Citrine uptake by tobacco BY-2 cells in the presence of 90 μM dTat-Sar-EED5.
Movie S3. Citrine uptake by tobacco BY-2 cells in the presence of 90 μM dTat as a control.

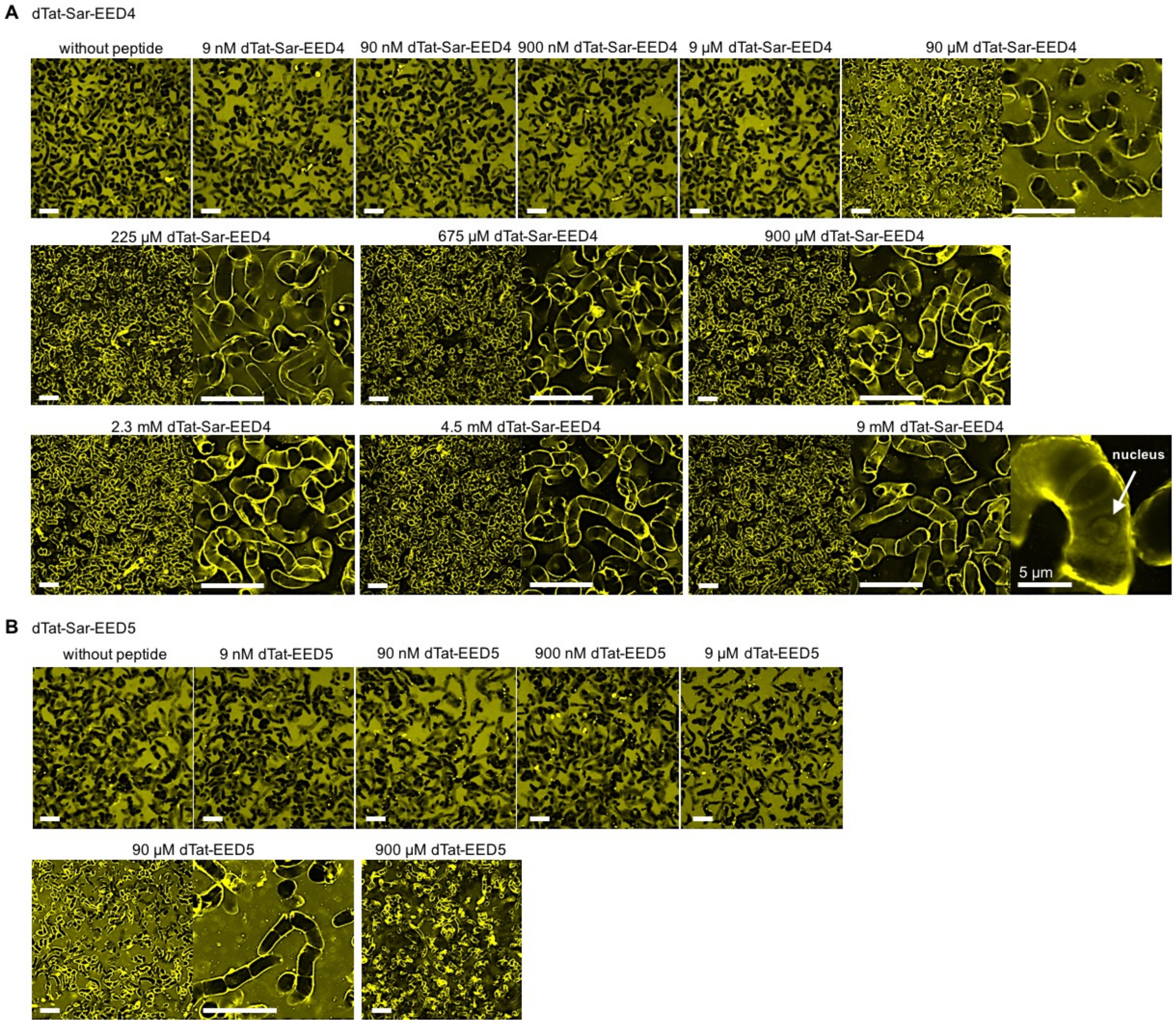


**Figure S1. Citrine internalization into tobacco BY-2 cells mediated by dTat-Sar-EED4 and dTat-Sar-EED5.** (**A**) Confocal microscopic images showing cells treated for 1 h with Citrine alone and in combination with dTat-Sar-EED4 at various concentrations. Enlarged images of 225, 675, and 900 µM (second row) and 2.3, 4.5, and 9 mM dTat-Sar-EED4 (third row) show the localization of Citrine. Scale bars indicate 50 µm unless otherwise stated. The white arrow shows a nucleus with detectable Citrine localization. (**B**) Confocal microscopic images showing cells treated for 1 h with Citrine alone and in combination with dTat-Sar-EED5 at various concentrations. An enlarged image of 90 µM dTat-Sar-EED5 shows the localization of Citrine. Scale bars indicate 50 µm.

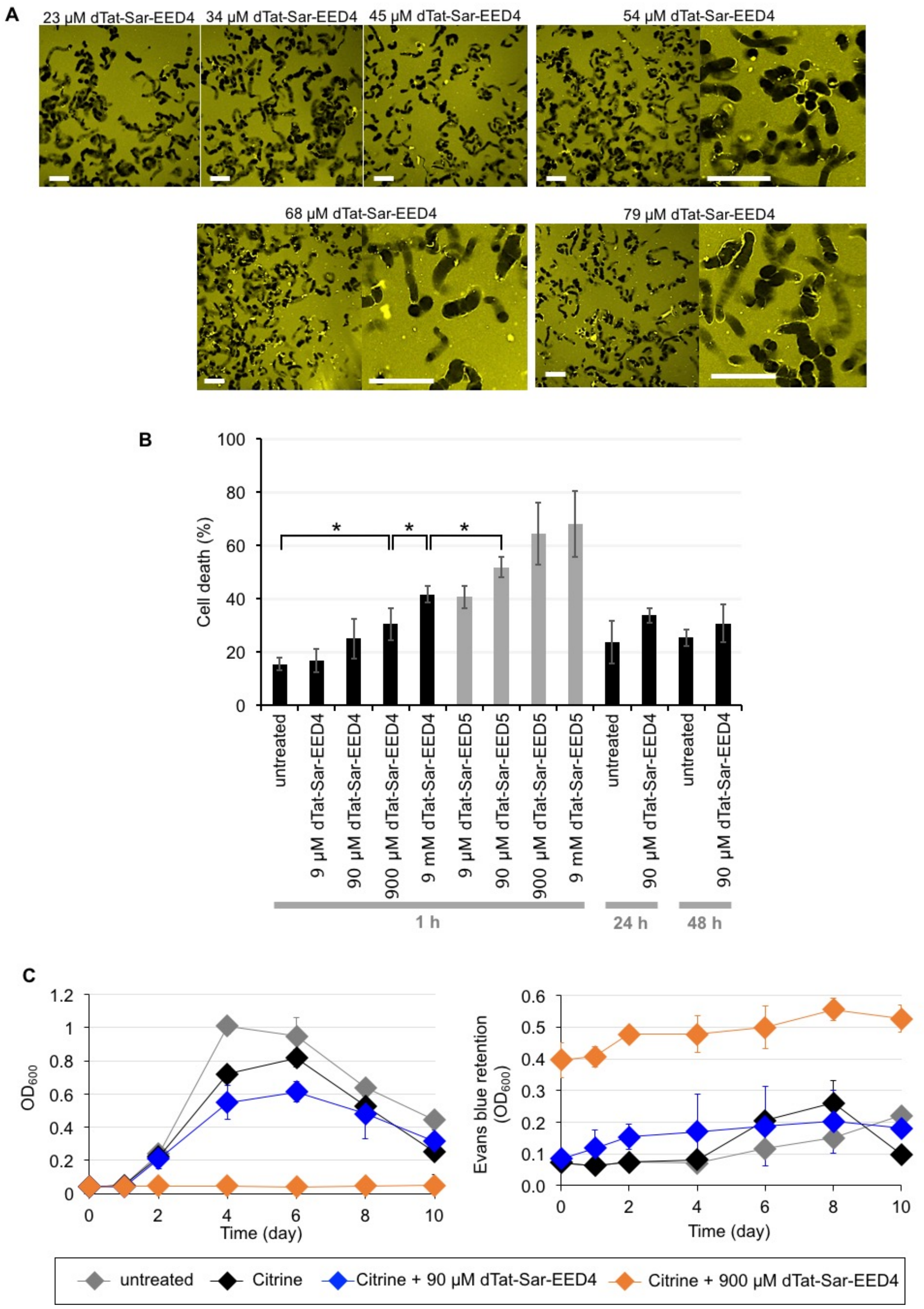


**Figure S2. Cytotoxicity effect of dTat-Sar-EED peptides on BY-2 cells.** (**A**) Confocal microscopic images showing cells treated for 1 h with Citrine in combination with dTat-Sar-EED4 at various concentrations. Enlarged images of 54, 68, and 79 µM dTat-Sar-EED4 show the localization of Citrine. Scale bars indicate 50 µm. (**B**) Cell death ratios estimated based on the Evans blue retention assay by using spectrophotometry. The cell death ratios are based on Evans blue retention. Data represent the mean values ± s.d. ($n$ = 3, *P < 0.05). (**C**) The time course of cell growth (left) and cell death ratio (right) monitored by spectrophotometry. Data represent the mean values ± s.d. ($n$ = 3).

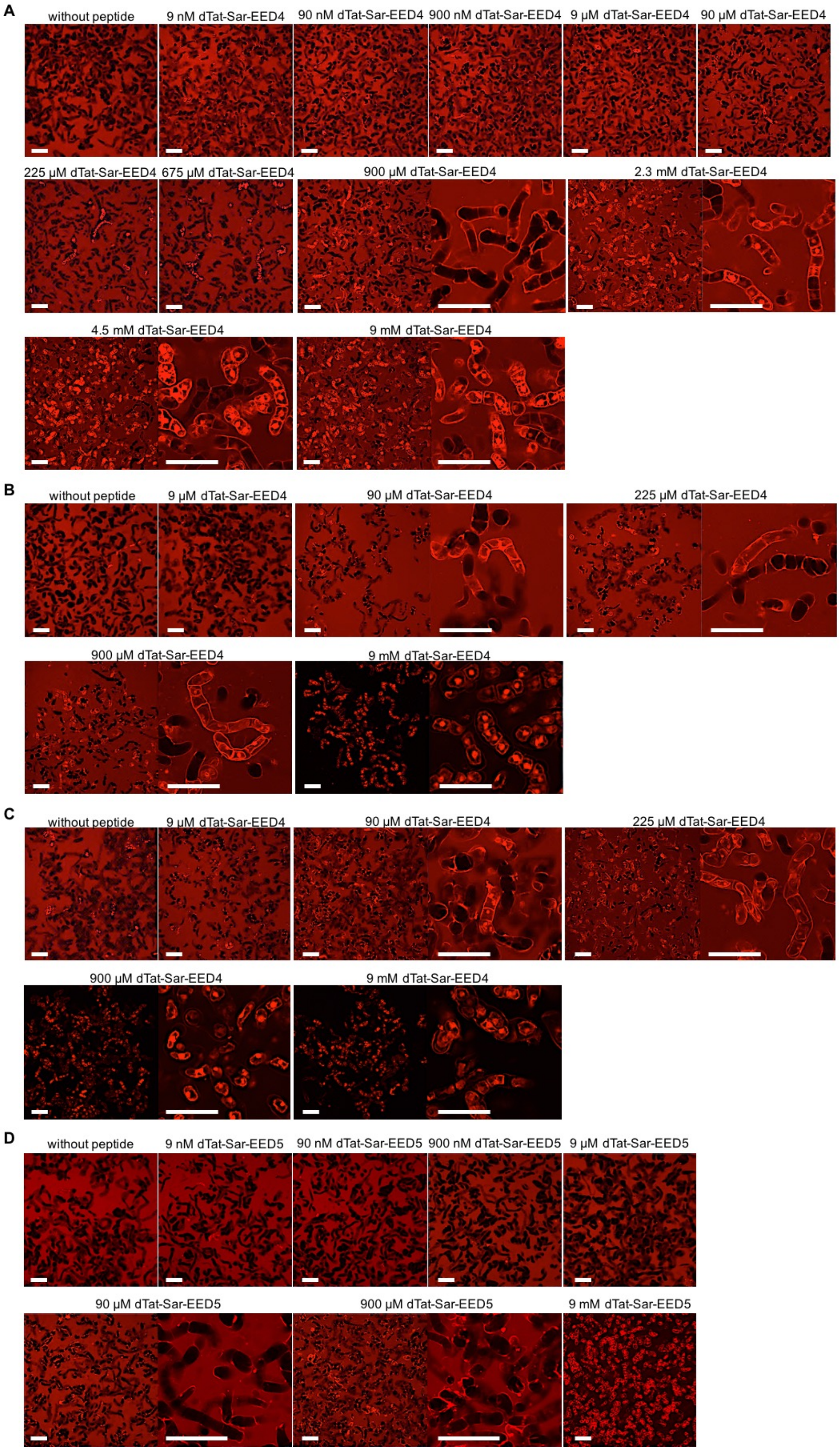


**Figure S3. Dextran internalization into tobacco BY-2 cells mediated by the dTat-Sar-EED peptides.** Confocal microscopic images showing cells treated for 1 h (**A**); 24 h (**B**); and 48 h (**C**); with dextran (TR-Dex70) alone and in combination with dTat-Sar-EED4 at various concentrations. (D) Confocal microscopic images showing cells treated for 1 h with TR-Dex70 alone and in combination with dTat-Sar-EED5 at various concentrations. Scale bars indicate 50 µm.

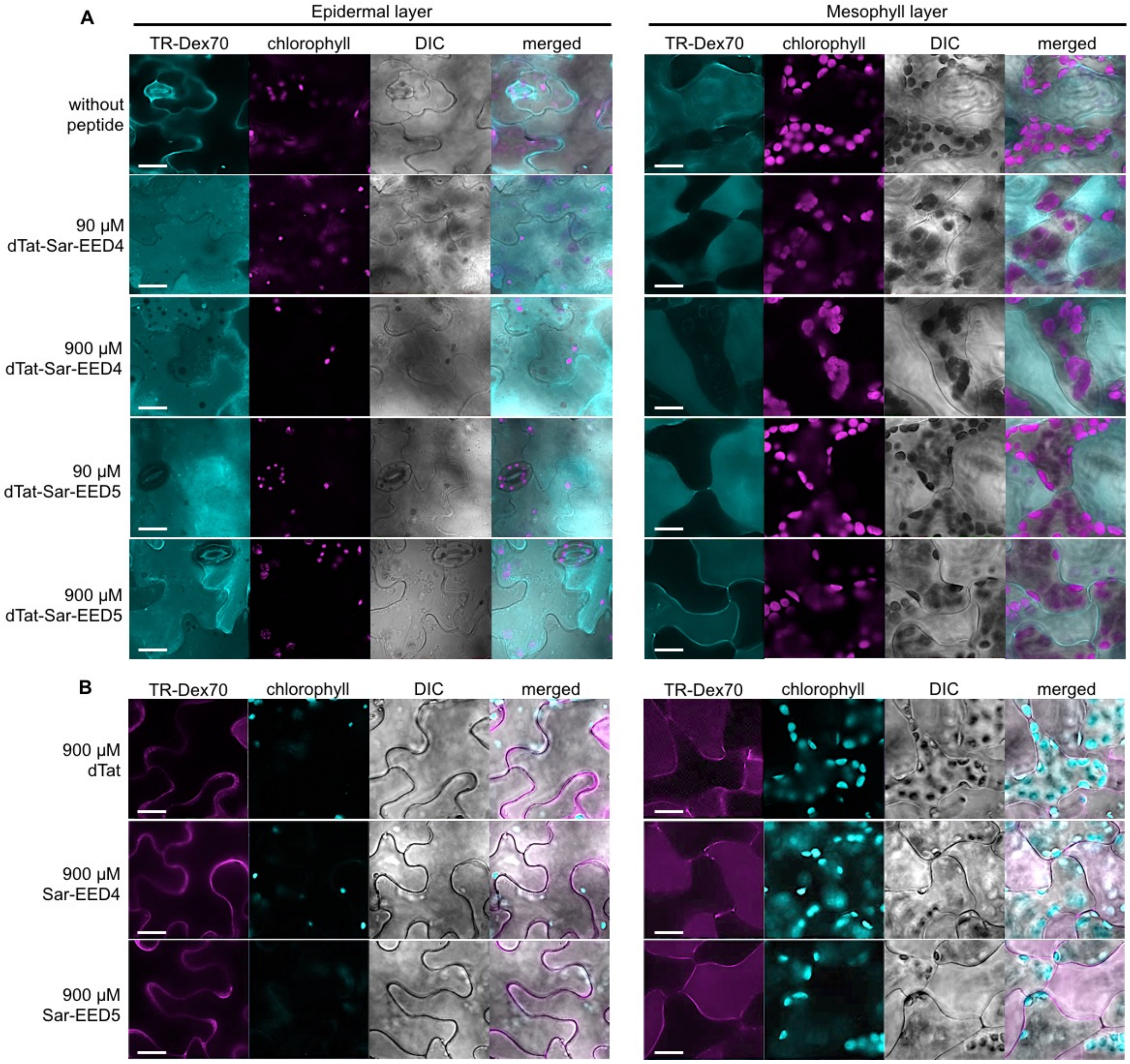


**Figure S4. Dextran internalization into *Arabidopsis thaliana* leaf epidermal and mesophyll cells mediated by the dTat-Sar-EED peptides.** (**A**) Confocal microscopic images showing Arabidopsis leaf cells treated for 1 h with dextran (TR-Dex70) alone and in combination with dTat-Sar-EED4 or dTat-Sar-EED5 at 90 or 900 µM. Scale bars indicate 10 µm. (**B**) TR-Dex70 internalization into Arabidopsis leaf epidermal and mesophyll cells mediated by 900 µM of dTat, Sar-EED4, or Sar-EED5 as controls. Scale bars indicate 10 µm.

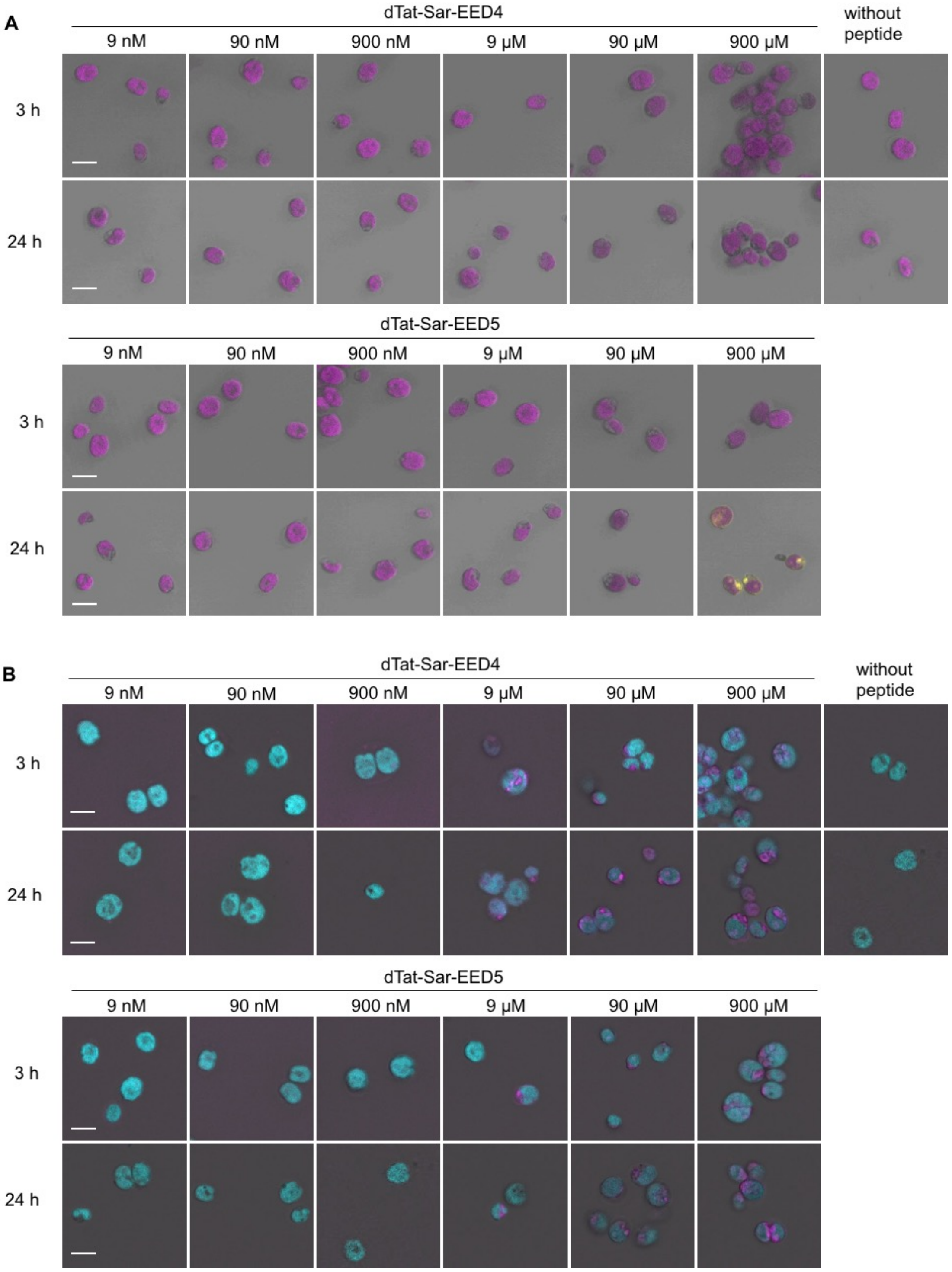


**Figure S5. Cargo internalization into *Chlamydomonas reinhardtii* cells mediated by the dTat-Sar-EED peptides.** (**A**) Confocal microscopic images showing *C. reinhardtii* cells treated for 3 h or 24 h with Citrine alone and in combination with each dTat-Sar-EED peptide at various concentrations. Yellow and magenta signals are Citrine and chlorophyll, respectively. Scale bars indicate 10 µm. (**B**) Dextran (TR-Dex70) internalization into *C. reinhardtii* cells mediated by the dTat-Sar-EED peptides. Magenta and cyan signals are TR-Dex70 and chlorophyll, respectively. Scale bars indicate 10 µm.

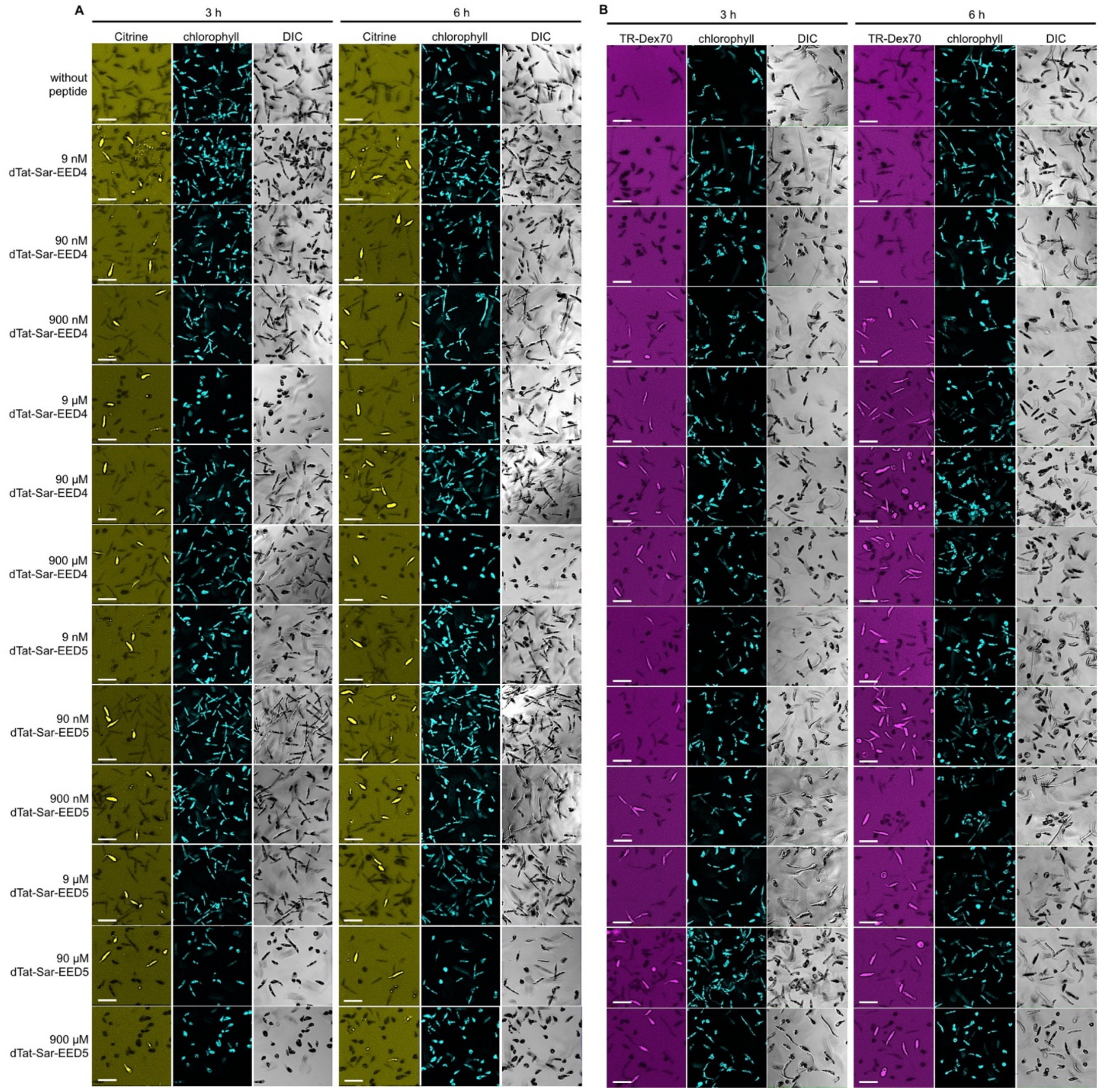


**Figure S6. Cargo internalization into *Euglena gracilis* cells mediated by the dTat-Sar-EED peptides.** (**A**) Confocal microscopic images showing *E. gracilis* cells treated for 3 h or 6 h with Citrine alone and in combination with each dTat-Sar-EED peptide at various concentrations. Scale bars indicate 50 µm. (**B**) Confocal microscopic images showing *E. gracilis* cells treated for 3 h or 6 h with dextran (TR-Dex70) alone and in combination with each dTat-Sar-EED peptide at various concentrations. Scale bars indicate 50 µm.

**Supplementary Movies**

**Movie S1.** Time-lapse imaging of Citrine uptake by tobacco BY-2 cells in the presence of 90 μM dTat-Sar-EED4.

**Movie S2.** Time-lapse imaging of Citrine uptake by tobacco BY-2 cells in the presence of 90 μM dTat-Sar-EED5.

**Movie S3.** Time-lapse imaging of Citrine uptake by tobacco BY-2 cells in the presence of 90 μM dTat as a control.
